# 3D Endothelium-on-a-Chip Reveals Cx43/LDHA Axis as a Driver of Endothelial Barrier Dysfunction in Diabetic Kidney Disease

**DOI:** 10.1101/2025.05.19.654627

**Authors:** Peter Lialios, Yoontae Kim, Isabella Trewhella, Komuraiah Myakala, Eleni Hughes, Xiaoxin Wang, Moshe Levi, Stella Alimperti

## Abstract

Endothelial dysfunction is a key pathological feature of diabetic kidney disease (DKD), characterized by increased vascular leakiness and altered metabolic signaling. In this study, we investigated how diabetic conditions affect endothelial barrier integrity and identified the molecular mechanisms contributing to this dysfunction. Using a 3D microfluidic model that recapitulates in vivo vascular architecture and flow, we demonstrated that high glucose (HG) and high fat (HF) conditions significantly impair endothelial barrier function, as demonstrated by increased dextran permeability and loss of VE-cadherin from the cellular membrane site. Metabolomic profiling and functional assays revealed a shift toward glycolysis, marked by elevated lactate levels and upregulation of lactate dehydrogenase A (LDHA), which contributed to barrier disruption. Pharmacological inhibition of LDHA effectively restored barrier function, underscoring the pathogenic role of glycolytic reprogramming. Transcriptomic analyses of mouse and human DKD datasets further identified connexin 43 (Cx43) as a candidate mediator of this dysfunction. Cx43 expression was progressively deregulated in diabetic mouse kidneys and across multiple cell types in human DKD samples. In vitro, Cx43 overexpression in endothelial cells enhanced glycolytic flux, suppressed oxidative metabolism, disrupted VE-cadherin localization, and promoted angiogenic sprouting. Collectively, our findings establish a mechanistic link between Cx43-driven metabolic reprogramming and endothelial barrier dysfunction in DKD, highlighting Cx43 as a potential therapeutic target for preserving vascular integrity in diabetic conditions.

## 1. INTRODUCTION

Diabetes affects over 37 million Americans, with type 2 diabetes (T2D) comprising the vast majority (90–95%) (1). A frequent and serious complication of T2D is diabetic kidney disease (DKD), a major cause of chronic kidney disease and eventual kidney failure(2–8). Although treatments like SGLT2 inhibitors can slow its progression (9–16), a thorough understanding of the molecular mechanisms driving DKD and glomerular scarring remains essential. Research indicates that damage to the glomerular endothelial cells (GECs), impaired microvascular barrier function, and increased permeability of kidney blood vessels are key factors in diabetic kidney pathophysiology (17–19). Albeit subtle, sustained leakage in the glomeruli can trigger chronic kidney inflammation and tissue damage (18, 20–23). Endothelial dysfunction often encompasses changes in the mesangium, microangiopathy, and enlarged glomerular capillaries (24–27). Considering the crucial role of glomerular microvasculature in both healthy and diseased kidneys, it is important to shed more light upon the regulation of glomerular microvascular barrier integrity to potentially uncover new therapeutic targets for DKD-related glomerulosclerosis.

In DKD, endothelial dysfunction is closely linked to aberrant glycolytic activity, particularly through the upregulation of lactate dehydrogenase A (LDHA), a key enzyme that catalyzes the conversion of pyruvate to lactate during glycolysis(28). Hyperglycemia drives increased glycolytic flux in endothelial cells, resulting in excessive lactate production and metabolic imbalance(28–30). LDHA plays a pivotal role in sustaining this glycolytic phenotype by maintaining NAD^+^ levels necessary for continuous glycolytic throughput(29, 31)^**11**,**14**^. In this context, elevated LDHA expression contributes to pathological angiogenesis and endothelial activation, namely hallmarks of disease progression (29, 30). Enhanced lactate production not only reflects a shift toward aerobic glycolysis (Warburg-like effect) but also promotes an inflammatory microenvironment that exacerbates glomerular injury (32–35). The inhibition of LDHA has been shown to reduce lactate accumulation, normalize endothelial metabolism, and attenuate inflammation and fibrosis in diabetic kidneys (32, 33). Furthermore, targeting LDHA may reverse endothelial-to-mesenchymal transition (EndoMT) and preserve the integrity of the glomerular filtration barrier, highlighting its potential as a therapeutic target in DKD (36, 37).

In parallel with metabolic reprogramming, the dysregulation of endothelial junctional complexes further compromises glomerular barrier integrity in DKD. Glomerular permselectivity and albuminuria are associated with disruption of GECs barrier via alterations in expression of key intracellular molecules, such as VE-Cadherin and ZO-1(38–45). Changes in cell adhesion can compromise the endothelial barrier function, contributing to glomerulosclerosis (46–52). Furthermore, connexins, namely proteins that form gap junctions and hemichannels allowing intercellular interactions, have been implicated in regulating glomerular function, vascular tone, and tubular integrity in the kidney (53–56). Of these connexins, Cx43 is highly expressed in GECs and regulates glomerular endothelial barrier function (57–60). In DKD, Cx43 expressions are altered in endothelial cells, while Cx43 has been shown to modulate processes such as infiltration, fibrosis, and epithelial-mesenchymal transition through pathways including SIRT1-HIF-1α (54, 61). In addition, Cx43 dysregulation in GECs disrupts NO production, glycocalyx degradation, and exacerbating leakiness and oxidative stress (56, 62). Animal studies have shown that Cx43 inhibition reduces albuminuria, glomerulosclerosis, and tubular damage (56, 63, 64).

Although the contributions of metabolic factors such as glucose and lipid accumulation to DKD are well-documented (65), their regulatory role on glomerular endothelial barrier integrity— particularly through cell adhesion molecules like Cx43—remains poorly understood. To address this gap, we investigated the interplay between metabolic dysregulation and structural dysfunction in the diabetic glomerular endothelium. In vivo studies revealed a significant upregulation of both LDHA and Cx43 protein levels in diabetic mice. Supporting this, single-nucleus RNA sequencing (snRNA-seq) data from diabetic mouse kidneys demonstrated increased Cx43 expression in glomerular endothelial cells, aligning with snRNA-seq datasets from human DKD (66). To further explore the functional role of Cx43 in regulating endothelial barrier integrity and glycolysis, we developed a 3D microfluidic platform that recapitulates the glomerular microvascular environment. This model enables the study of rapid, physiologically relevant responses in endothelial cells, overcoming the limitations of conventional 2D cultures and avoiding the complexity of in vivo models. Exposure of glomerular endothelial cells to high glucose and fatty acid conditions led to increased endothelial permeability and elevated LDHA expression. Pharmacological inhibition of LDHA restored barrier function, suggesting that hyperglycemia compromises endothelial integrity via LDHA-mediated glycolytic dysregulation. This loss of barrier function was accompanied by reduced VE-Cadherin expression and increased Cx43 protein levels, indicating impaired cell–cell adhesion. Furthermore, Cx43 overexpression in endothelial cells resulted in heightened barrier leakiness and upregulation of LDHA, supporting a mechanistic link between Cx43 activity and metabolic reprogramming under diabetic conditions. Collectively, our findings highlight a novel Cx43–LDHA axis driving glomerular endothelial dysfunction in DKD. This study sheds light on the glycolytic regulation of endothelial integrity and proposes a potential therapeutic target to preserve glomerular function by modulating the interaction between Cx43 and metabolic pathways in response to diabetic stress.

## 2. MATERIALS AND METHODS

### Cell Culture

Human umbilical vein endothelial cells (HUVECs) (Lonza) at passage 2 to 4 and glomerular endothelial cells (GECs) (ScienCell) at passage 3-4 were cultured in supplemented Endothelial Growth Medium-2 (EGM-2) (Lonza) and Endothelial Cell Medium (ECM) (ScienCell), respectively. For diabetic conditions, cells were exposed to; high glucose (HG): 25 mM D-glucose, high fat (HF): media containing 4.5 mg/L cholesterol, 10 mg/L fatty acids, and 2 mg/L sodium acetate (Sigma: L5146), and their combination (HG+HF) for 48 h, compared to normal glucose (NG: 2.7mM) conditions. For blocking glycolysis, cells were exposed to LDHA inhibitor (GSK2837808; 5 μM, MedChemExpress) in presence of the HG+HF treatment. To obtain Cx43 overexpression (Cx43over), the cells were transduced sequentially with a blasticidin-resistant vector encoding dCas9-VP64-Blast (transcriptional activator) and a hygromycin-resistant vector encoding and lenti MS2-P65-HSF1-Hygro (co-activator complex) and pLenti-sgRNA(MS2)-zeo vector (GJA1 SAM guide (GGGACATGAACGCCTCTACT)).

### Fabrication of Microfluidic Device and Measurements of Endothelial Barrier Function

Microfluidic devices were constructed as shown previously (67–69). Briefly, polydimethylsiloxane (PDMS; Sylgard 184, Dow-Corning; Krayden) devices were created from 3D printed scaffolds and mounted on an inverted cast to form cylindrical channels with a 160 μm diameter in a hydrogel. These devices were then treated with 0.01% (v/v) poly-L-lysine (PLL; Sigma) and 0.5% (v/v) glutaraldehyde (Sigma), followed by an overnight wash in diH2O. To form perfusable microvascular channels, 160 µm diameter steel acupuncture needles (Seirin, Kyoto, Japan) were inserted. Subsequently, a hydrogel mixture of 3mg/mL Collagen I (BD) was introduced and incubated overnight as shown previously (67–69). After removing the needles, endothelial cells (HUVECs or GECs) were seeded into the channels at a concentration of 1 × 10^6^ cells/ml. To facilitate cell attachment, devices were inverted for 2 min and then flipped upright for another 2 min. Non-adherent cells were removed by scraping and flushing with fresh media, and the devices were cultured on a platform rocker (BenchRocker BR2000) at 37 °C in a CO2 incubator for 24 h under the corresponding conditions. To evaluate endothelial permeability within the microfluidic system, fluorescent dextran (70 kDa Texas Red, Thermo Fisher) was added to the perfusion medium at a concentration of 12.5 µg/mL. The diffusion of dextran was monitored in real-time using a confocal microscope (Evident Olympus) with a 10× objective. Time-lapse images were analyzed by calculating the average fluorescence intensity in regions adjacent to the endothelial layer across sequential frames. The rate of intensity change over time was determined through linear regression for each region. This rate, along with the average intensity (I) and capillary radius (r), was used to calculate the diffusive permeability coefficient (P_d_) using the appropriate 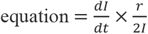. Measurement variability was quantified by calculating the standard deviation of permeability values across different regions, following previously established methods (67–69)

### Immunofluorescence Staining

Cells were fixed with 4% v/v paraformaldehyde (PFA) (Sigma) in PBS (pH 7.4) for 30 minutes at room temperature (RT), followed by washing twice with Phosphate Buffered Saline (PBS). Next, the cells were permeabilized with 0.1% Triton X-100 in PBS for 30 min and blocked with a solution containing 0.01% Triton X-100, 5% goat serum (Sigma) in PBS overnight at 4°C. Next, samples were incubated with anti-human VE-Cadherin primary antibodies (1:1000 dilution; #2500, Cell Signaling Technology) in blocking solution overnight at 4°C. The cells were washed twice, followed by the application of Alexa Fluor 568-conjugated goat-anti-rabbit IgG secondary antibody (1:100 dilution; Thermo Fisher) overnight at 4°C. Nuclei were counterstained with DAPI (1:300 dilution; Thermo Fisher) in blocking solution for 10 minutes. F-actin was visualized using Alexa Fluor 647–conjugated phalloidin (1:200 dilution; Thermo Fisher), and an additional F-actin channel was labeled using Alexa Fluor 488–phalloidin (1:200 dilution; Thermo Fisher), where indicated. Confocal imaging was performed using a Nikon CSU-W1 SoRa Spinning Disk Microscope. Z-stack projections, channel merging, and cross-sectional analysis were generated using ImageJ All western blots were quantified by Image J according to standard procedures (70).

### Metabolomics and snRNAseq

Whole homogenized protein extracts from Six-week-old male db/m and db/db mice (BLKS/J) were collected and injected into a Waters Xevo G2 LC-MS for untargeted profiling of metabolites (Georgetown University, DC, USA). Data were collected in both electrospray ionization (ESI) positive (POS) and negative (NEG) modes. Analysis of Cx43 expression levels in human diabetic kidney disease (DKD) was conducted using the Kidney Interactive Transcriptomics (KIT) databas (https://humphreyslab.com/SingleCell/)

### Animal Studies

Six-week-old male db/m and db/db mice (BLKS/J) were obtained from the Jackson Laboratories. They were maintained on a 12-h light/12-h dark cycle. They were on (*1*) a regular chow diet, (*2*) INT-767 (30 mg/kg body wt/d), (*3*) INT-747 (20 mg/kg body wt/d), or (*4*) INT-777 (30 mg/kg body wt/d) admixed with chow for 2 weeks.

### Western Blot Analysis

Protein lysates were extracted from HUVECs cultured under normal glucose (NG; 5.5 mM), high glucose (HG; 25 mM), high fat (HF; 4.5 mg/L cholesterol, 10 mg/L fatty acids), and HG+HF conditions using RIPA lysis buffer (Thermo Fisher) supplemented with protease and phosphatase inhibitors (Cell Signaling Technology). For the in vivo studies, cortical homogenate protein was obtained from mice. For both in vitro and in vivo studies, protein concentrations were quantified via bicinchoninic acid (BCA) assay (Pierce™), and 20 µg of lysate per sample was resolved on 4% to 12% gradient SDS–PAGE gels (Thermo Fisher). Proteins were transferred to nitrocellulose membranes (Thermo Fisher) and blocked with 5% w/v bovine serum albumin (BSA) (Sigma) in Tris-buffered saline with 0.1% v/v Tween 20 Detergent (TBST) (Thermo Fisher) for 1 h. Membranes were probed overnight at 4°C with the following primary antibodies: rabbit anti-human LDHA (1:1,000 dilution; #3582, Cell Signaling Technology), rabbit anti-human Cx43 (1:1,000 dilution; #3512, Cell Signaling Technology), and rabbit anti-human β-actin (1:5,000 dilution; #4970, Cell Signaling Technology). The antibodies were diluted and incubated in 5 % w/v BSA in TBST (Thermo Fisher) overnight in 4 °C. Proteins of interest were detected with horseradish peroxidase-conjugated (HRP-linked) anti-rabbit IgG secondary antibodies (1:1,000 dilution; #7074, Cell Signaling Technology) and protein bands were visualized with chemiluminescent substrate Pico PLUS (Thermo Fisher) and Azure 300 Imaging System (Azure Biosystems), according to manufacturer instructions. All western blots were quantified by ImageJ according to standard procedures (70).

### Statistical Analysis

Statistical analysis was conducted using GraphPad Prism 10. Sample sizes for each experimental group are provided in the figure legends (N). All in vitro experiments were performed in triplicate and independently repeated three times. Data was assessed for normality, and no significant variation between groups was detected. For comparisons between two groups, a two-tailed Student’s *t*-test was applied. For experiments involving multiple time points or treatment conditions, one-way or two-way ANOVA was used, followed by Tukey’s post hoc test where appropriate. For in vivo studies, the unit of measurement was the individual animal. A two-way ANOVA was used to assess differences between groups. Sex was included as a biological variable in the experimental design. A priori power analysis, assuming a type I error rate of 5% and a statistical power of 90%, indicated that a sample size of N = 5 animals per group was sufficient to detect statistically significant differences. All in vivo experiments were conducted and analyzed in a blind manner to ensure unbiased interpretation. Results are presented as mean ± standard deviation (SD), and *p* < 0.05 was considered statistically significant.

## 3. RESULTS

### 3.1 Diabetic conditions impair endothelial barrier function

Diabetic db/db mice demonstrate an increase in serum glucose and lipid levels compared to nondiabetic db/m mice (71–73). The metabolic consequences of glucose and lipid dysregulation in obesity and diabetes mellitus result in endothelial cell dysfunction with delicate and leaky blood vessels (74–79). Similar to those findings, we pursued studies to evaluate how hyperglycemic conditions alter barrier function. We developed and utilized a 3D microfluidic platform that mimics blood vessel structure and flow forces in vivo (80, 81). The 3D microfluidic device **(Fig 1A)** is fabricated with a 3D printed scaffold in which either human glomerular endothelial cells (GECs) (82, 83) and HUVECs (Lonza) have been seeded as shown previously (67–69). The microfluidic device was mounted on an inverted cast to form cylindrical channels (160 µm diameter) in a hydrogel. Upon seeding the endothelial cells, a perfusable microvessel was formed, as described in **Fig 1B,C** and our publications (67, 69, 84). The fluorescent assay assessed Endothelial barrier function (Red labeled 70 kDa dextran) as shown in **Fig 1D** and our recent publications (67, 69, 84). Specifically, our data showed that upon the presence of high glucose (HG: 25mM) and/or high fat (HF) conditions (media: containing 4.5 mg/l cholesterol, 10mg/l fatty acids, 25 mg/l polyoxyethylene sorbitan monooleate and 2mg/l acetate), the endothelial leakiness is increased by 10-times compared to cells under normal glucose levels (NG: 2.7mM) (**Fig 1D**). Finally, we examined the role of VE-Cadherin, a critical intracellular adhesion molecule whose altered expression or localization is associated with impaired endothelial barrier integrity(38–45). Consistent with previous studies, our immunostaining results revealed that exposure to HG+HF conditions led to the loss of VE-Cadherin from endothelial junctions. Specifically, the intensity of junctional VE-Cadherin staining was reduced by ~2.2-fold (p < 0.05) relative to NG controls, indicating that diabetic conditions significantly compromise endothelial barrier function (**Fig. 1E**). These findings collectively demonstrate that diabetic conditions impair endothelial barrier integrity by disrupting VE-Cadherin localization, highlighting a potential mechanistic link between metabolic dysfunction and vascular barrier dysregulation.

**Figure 1.**
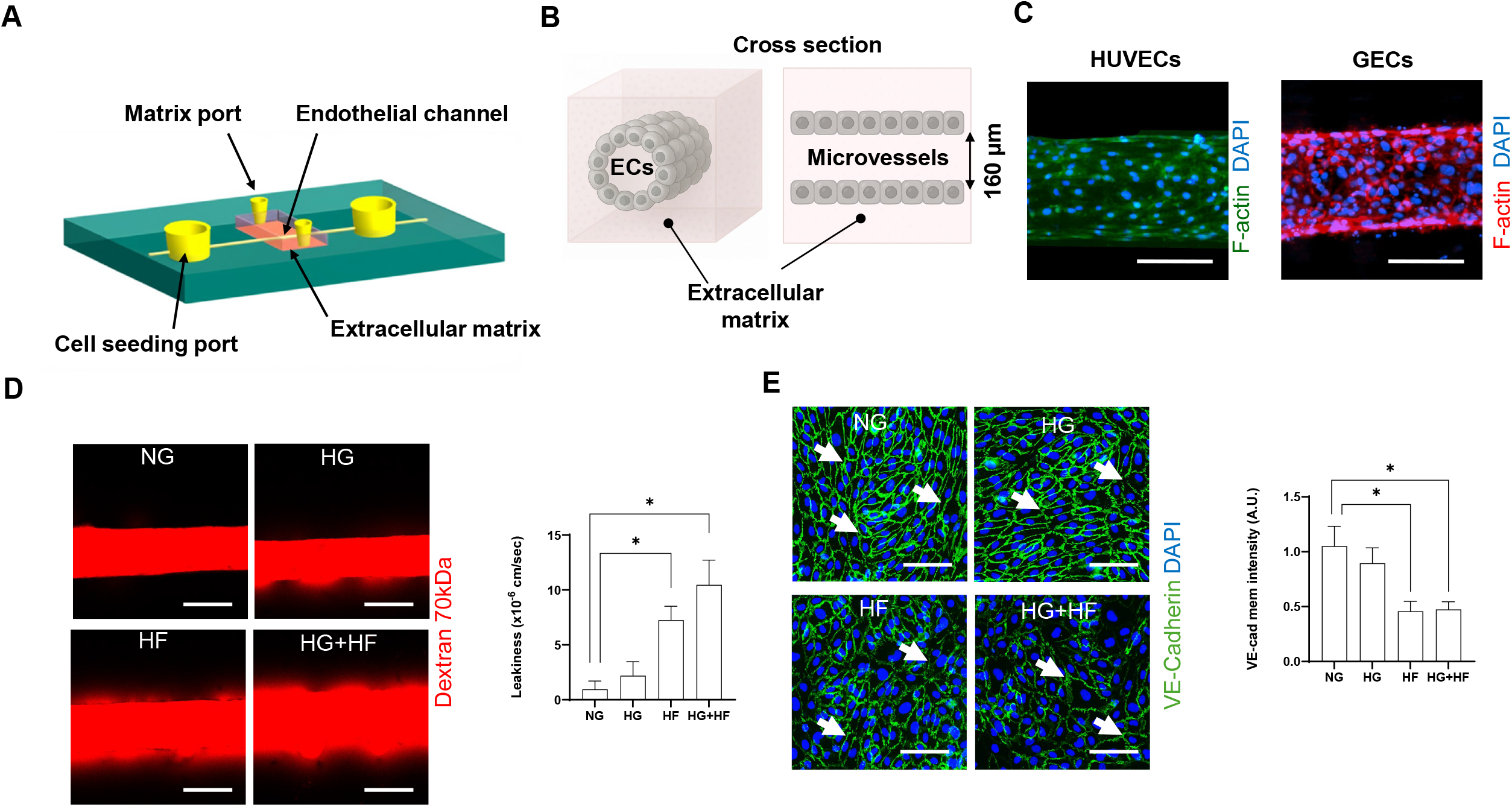
Design and application of a 3D microvascular microfluidic system to assess endothelial barrier function under normal and diabetic-like metabolic conditions. **(A)** Schematic illustration of the 3D vascularized microfluidic platform, featuring endothelial channels, matrix ports, and cell seeding ports within an extracellular matrix scaffold. **(B)** Cross-sectional diagram of the engineered microvessel structure showing a lumen surrounded by endothelial cells (ECs) embedded in extracellular matrix. **(C)** Representative confocal images of engineered microvessels formed using HUVECs and glomerular endothelial cells (GECs) embedded in a 3D extracellular matrix. Cells were stained for F-actin (green/red) and nuclei (DAPI, blue). Scale bars = 100 µm. **(D)** Representative images showing the diffusion of 70 kDa fluorescent dextran (red) across endothelial microvessels under NG, HG, HF, and HG+HF conditions. Scale bars = 100 µm. Quantification of endothelial leakiness (measured as diffusive permeability coefficient, Pd × 10−6 cm/sec) reveals significantly increased permeability under HF and HG+HF treatments (*p < 0.05). **(E)** Immunofluorescence staining of VE-Cadherin (green) and DAPI (blue) in endothelial monolayers under normal glucose (NG), high glucose (HG), high fat (HF), and combined HG+HF conditions. White arrows indicate VE-cadherin localization at cell-cell junctions. Quantification of VE-cadherin membrane intensity (A.U.) demonstrates a significant reduction in HG, HF, and HG+HF conditions compared to NG (p < 0.05).

### 3.2 Diabetic Conditions alter LDHA levels

To further elucidate the molecular mechanisms underlying endothelial dysfunction under diabetic conditions, we examined the metabolic profile of ECs. Similar to previous studies, elevated glucose levels in DKD have been linked to alterations in endothelial metabolism (29, 85–90). Our metabolomics analysis revealed significantly increased lactate levels in kidney tissue from db/db mice compared to db/m controls (*p* < 0.006) (**Fig 2A**), indicating a glycolytic metabolic shift. In addition, our data showed that high glucose/high fat (HG+HF) conditions enhance glycolysis via upregulation of lactate dehydrogenase A (LDHA) by ~3-times (p<0.01) compared to NG (**Fig 2B**). Moreover, pharmacologic inhibition of LDHA using GSK2837808 (5 µM) (91, 92) in the presence of HG+HF conditions, significantly reduced endothelial leakiness by 3-fold (p < 0.05) (**Fig. 2C**). These findings indicate that glycolytic reprogramming plays a critical role in endothelial barrier disruption under diabetic conditions, and that targeting key enzymes such as LDHA may offer a therapeutic strategy to restore vascular integrity.

**Figure 2.**
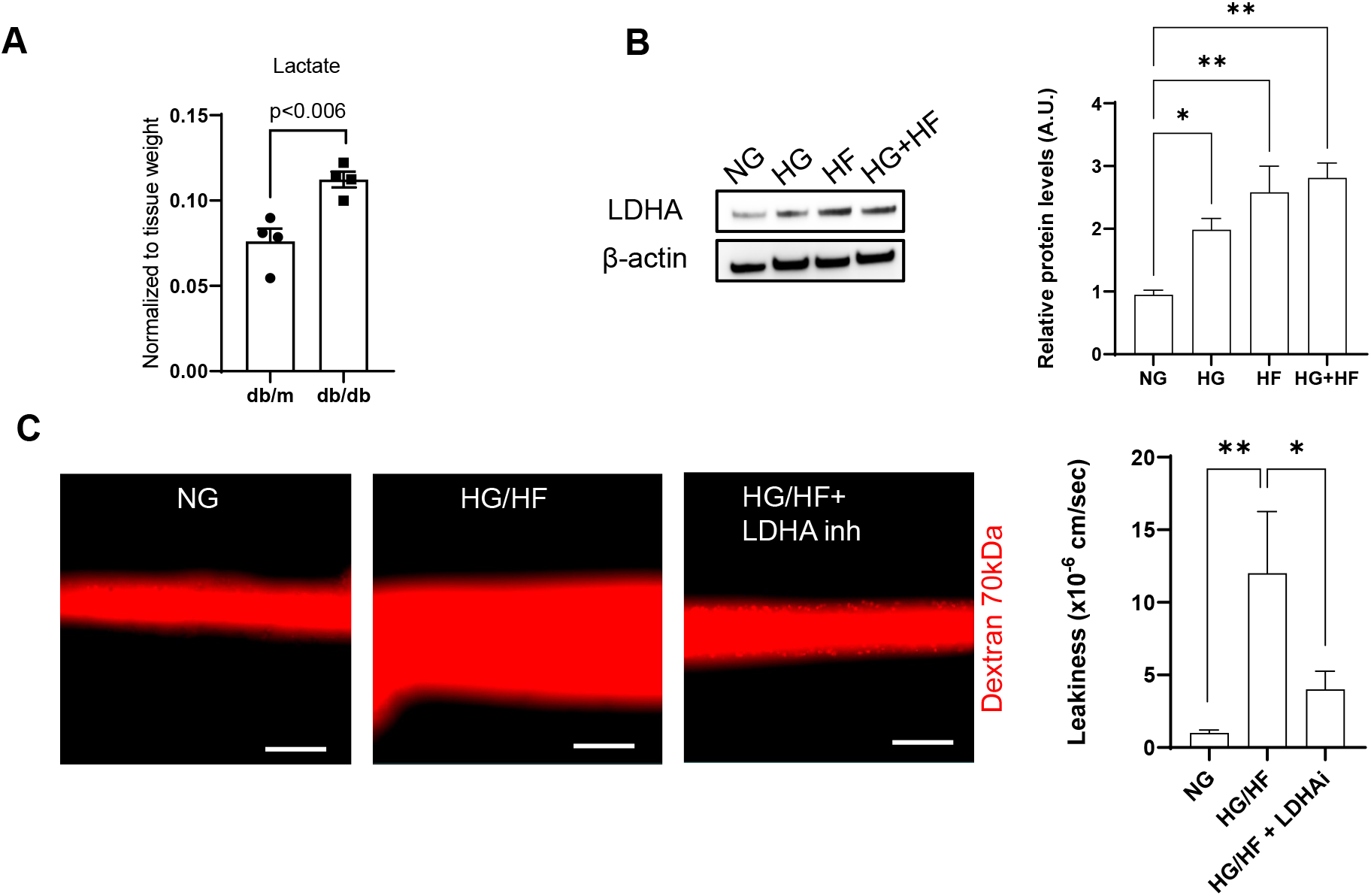
Elevated glucose and lipid levels promote metabolic reprogramming with glycolytic shifts, while blocking glycolysis restores endothelial barrier integrity. **(A)** Metabolomics analysis of kidney tissue from db/m and db/db mice demonstrates increased lactate levels in db/db mice (p < 0.006), normalized to tissue weight (n = 4 per group; mean ± SEM). **(B)** Western blot analysis of lactate dehydrogenase A (LDHA) and densitometric quantification of relative protein levels, with β-actin used as a loading control. High glucose and lipid treatment (HG+HF) significantly upregulates LDHA, indicating a shift toward glycolytic metabolism. (**p < 0.01 vs NG). **(C)** Analysis of leakiness measured as diffusive permeability coefficient (Pd × 10−6 cm/sec). Scale bars = 100 μm. HG+HF significantly impairs endothelial barrier function, causing higher permeability of 70 kDa dextran in the microfluidic device setup (**p < 0.01). Inhibition of LDHA rescues endothelial barrier function, as indicated by the decreased diffusion of 70 kDa dextran (*p < 0.05 vs HG+HF).

### 3.3 Diabetic Conditions Elevate Cx43 Expression in vivo and in vitro

Next, we investigated how diabetic conditions influence endothelial barrier integrity, focusing on the role of connexins. Connexins, which form gap junctions and hemichannels, are key mediators of intercellular communication and have been implicated in the regulation of glomerular function and vascular tone (53–56). Our data specifically highlights the upregulation of connexin 43 (Cx43) under diabetic conditions. Immunoblot analysis of kidney tissues from diabetic (db/db) and control (db/m) mice revealed a time-dependent increase in Cx43 protein expression. A significant upregulation—by ~1.8-fold (p < 0.001)—was detected at 26 weeks in diabetic mice (**Fig. 3A**). To extend these findings to human disease, we analyzed published single-nucleus RNA-sequencing and single-cell ATAC-seq datasets from human diabetic kidney disease (DKD) samples. These analyses revealed robust Cx43 expression across multiple cell types in DKD tissues (**Fig. 3B**) *(66, 93)*. To functionally validate these transcriptional trends, we performed RT-qPCR on endothelial cells exposed to metabolic stress. Gja1 (Cx43) mRNA expression was significantly upregulated under high glucose (HG), high fat (HF), and combined HG+HF conditions (**Fig. 3C**; p < 0.05). Consistently, immunoblotting confirmed increased Cx43 protein levels under HG (~**2.8-fold**), HF (~**6.6-fold**), and HG+HF (~**7.3-fold**) conditions (**Fig. 3D**; p < 0.05). These findings collectively indicate that Cx43 is progressively upregulated under diabetic conditions in both mouse models and human DKD, suggesting a potential role for Cx43 in mediating endothelial barrier dysfunction during disease progression.

**Figure 3.**
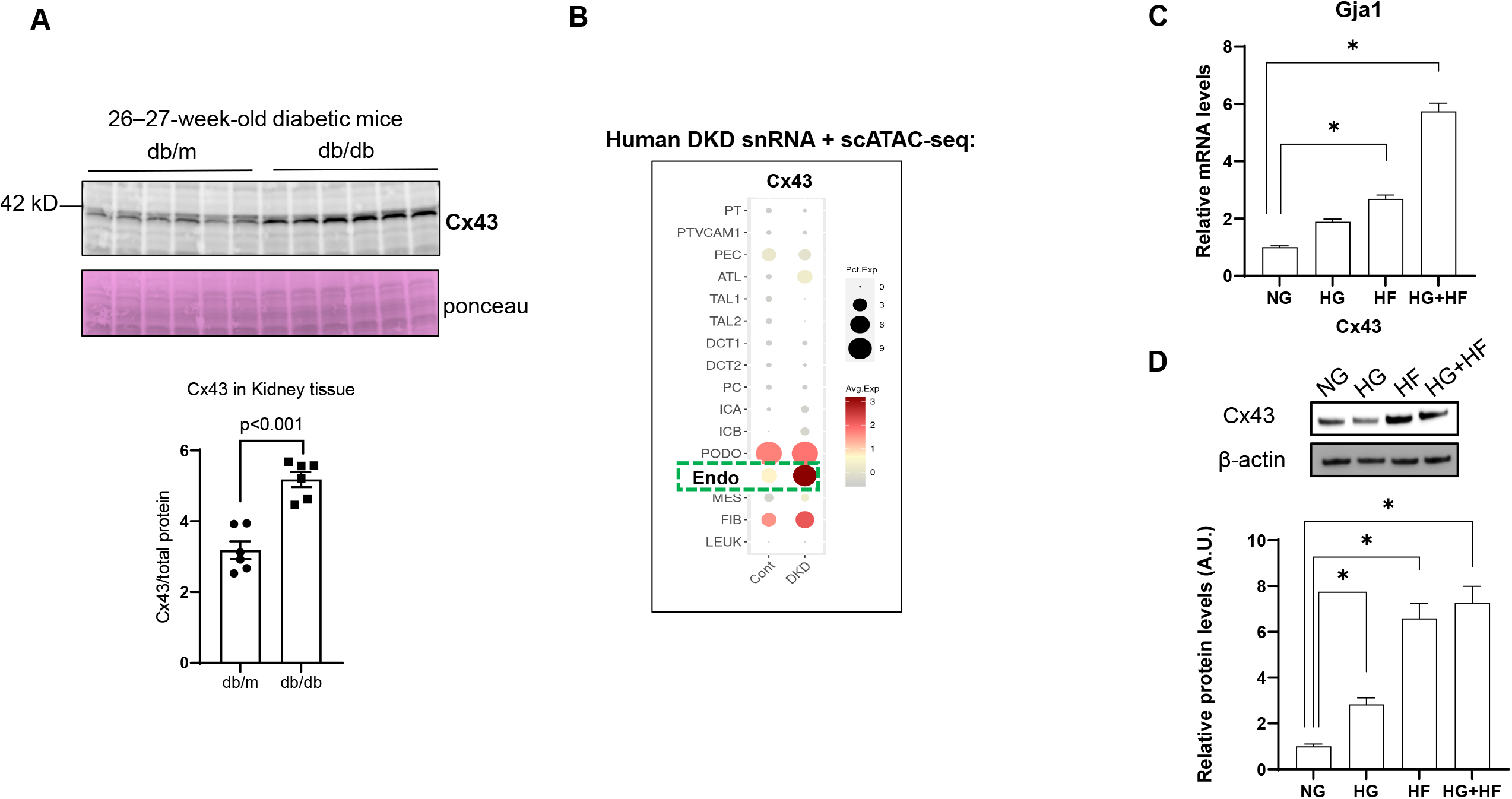
Cx43 expression is elevated in diabetic kidney disease across mouse models and human datasets. **(A)** Western blot analysis of Cx43 protein levels in kidney tissue from db/m (control) and db/db (diabetic) mice at 26–27 weeks of age. Ponceau staining is shown as a loading control. Quantification reveals significantly elevated Cx43 expression in db/db mice (p < 0.001; n = 5–6 per group; mean ± SEM). **(B)** Single-nucleus RNA-seq and single-cell ATAC-seq analysis of Cx43 (Gja1) expression in human diabetic kidney disease (DKD) datasets by Kidney Interactive Transcriptomics (KIT) (https://humphreyslab.com/SingleCell/). Dot size indicates percentage of cells expressing Cx43; red color intensity indicates average expression. Notably, endothelial cells (Endo, green dashed box) exhibit prominent expression in DKD. **(C)** Quantitative RT-PCR analysis of Gja1 (Cx43) mRNA levels in endothelial cells cultured under normal glucose (NG), high glucose (HG), high fat (HF), and combined HG+HF conditions. Cx43 transcript levels are significantly increased in HF and HG+HF conditions. (*p < 0.05; n=3; mean± SEM.**(D)** Immunoblotting and densitometric quantification of Cx43 protein under the same conditions as in (D) reveal elevated protein levels in HG and HF treatments (*p < 0.05; n = 3; mean ± SEM). β-actin serves as the loading control.

### 3.4 Cx43 Overexpression Promotes LDHA mediated Glycolytic Shift and Endothelial Dysfunction

Given that Cx43 is predominantly expressed in endothelial cells and plays a significant role in diabetic kidney disease (DKD)—not only by facilitating intercellular communication (37–40) but also by modulating key metabolic processes such as nitric oxide (NO) production (40, 42)— we sought to directly assess its role in endothelial barrier disruption and metabolic reprogramming. To this end, we overexpressed Cx43 (Cx43over) in endothelial cells (**Fig 4A**), as confirmed by immunoblot analysis. In addition, overexpression of Cx43 in endothelial cells results in an increase in LDHA protein expression by ~3.6-times (p<0.001). Finally, Cx43 overexpression led to significant endothelial barrier disruption, as evidenced by increased permeability to 70 kDa dextran in a 3D microfluidic assay (**Fig. 4B**). Quantitative analysis revealed a significantly ~11.7-times (p<0.01) increase in Cx43over endothelium compared to control (**Fig. 4C**; p < 0.001).. In parallel, sprouting assays demonstrated enhanced angiogenic activity under Cx43over conditions, with increased protrusions and a greater number of angiogenic extensions (**Fig. 4D**; p < 0.001). These findings suggest that Cx43 overexpression not only promotes a glycolytic metabolic phenotype but also compromises endothelial barrier integrity and enhances angiogenic behavior— linking Cx43 dysregulation to key pathogenic features of DKD.

**Figure 4.**
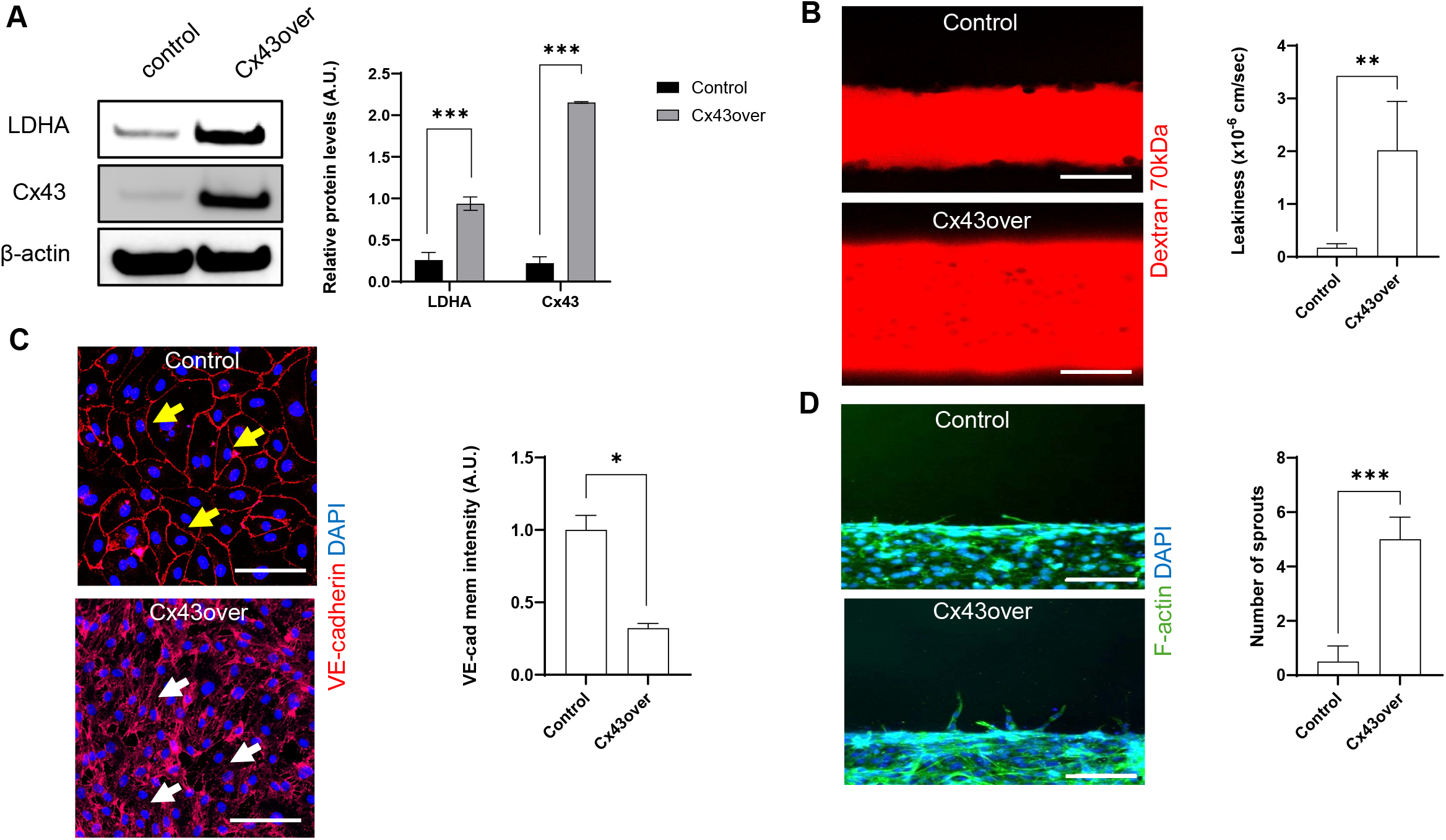
Cx43 overexpression disrupts endothelial barrier and junctional integrity, increases vascular permeability, and promotes angiogenic sprouting. **(A)** Representative immunoblot and corresponding densitometric analysis showing relative protein levels of Cx43 and lactate dehydrogenase A (LDHA) in Cx43 overexpressing (Cx43over) endothelial cells (ECs) compared to control ECs (***p < 0.001). β-actin was used as a loading control. **(B)** Representative images showing 70 kDa dextran (red) permeability across endothelial monolayers in microfluidic devices. (scale bars = 100 μm) and corresponding quantification of endothelial barrier function measured as diffusive permeability coefficient (Pd × 10−6 cm/sec). Significantly increased leakiness is observed in Cx43over cells compared to control (**p < 0.01). **(C)** Immunofluorescence staining of VE-cadherin (red) and nuclei (DAPI, blue) in control and Cx43over endothelial monolayers. Yellow arrows indicate intact junctional VE-cadherin localization in control cells, while white arrows highlight disrupted junctional VE-cadherin in Cx43over cells. Scale bars = 50 μm. **(D)** F-actin (green) and nuclear (DAPI, blue) staining showing endothelial sprouting in control and Cx43over conditions (scale bars = 100 μm) and corresponding quantification of endothelial sprout formation, showing significantly increased sprouting in Cx43over cells compared to control (***p < 0.001).

Overall, our results highlight Cx43 as a central regulator of endothelial dysfunction in diabetes and suggest that targeting Cx43 or its downstream metabolic pathways may offer new therapeutic avenues to preserve vascular integrity in diabetic kidney disease.

## 4. DISCUSSION

DKD has been characterized by profound vascular abnormalities, including endothelial dysfunction and increased glomerular permeability, which contribute to progressive renal damage and eventual kidney failure (17–27). In this study, we identified key molecular and metabolic mechanisms by which diabetic conditions disrupt endothelial barrier function. Specifically, we identified the central role of Cx43 in regulating glycolytic reprogramming and endothelial barrier function under diabetic stress conditions.

One potential challenge to elucidate these new mechanisms is that existing vivo models are too laborious for studying rapid structural and functional changes of the endothelium. Additionally, most of the existing conventional 2D *in vitro* models fail to recapitulate endothelial cell-mediated mechanosensitive interactions in healthy and disease states because they lack three-dimensional (3D) vascular biomimetic architecture and shear stress (94–97). Herein, we used a 3D microfluidic model of endothelium to elucidate how HG and HF conditions regulate barrier function. Our results demonstrated the increase in endothelial leakiness as shown in previous studies (74–79). This barrier disruption was closely associated with a loss of junctional VE-cadherin, a key component of adherens junctions that is essential for maintaining vascular integrity (38–45).Interestingly, our data further revealed that diabetic conditions promote a metabolic shift toward glycolysis in endothelial cells by upregulating the levels of LDHA. Importantly, the inhibition of LDHA under HG+HF conditions restored barrier function, suggesting a direct role for glycolytic flux in endothelial dysfunction.

Furthermore, connexins facilitate intercellular interactions, which regulate glomerular function, vascular tone, and tubular integrity in the kidney (53–56). Specifically, our data in conjunction with data extracted from Kidney Interactive Transcriptomics (66, 93) revealed that Cx43, Cx40, and Cx37 are majorly expressed in kidney endothelial cells. While Cx40 and Cx37 also play a role in regulating vascular tone and blood flow in the glomerulus, they are less involved in barrier function (60, 98). Of these connexins, Cx43 is highly expressed in GECs and regulates glomerular endothelial barrier function (57–60). In DKD, Cx43 expression is upregulated in endothelial cells. It has been shown that Cx43 promotes infiltration and interstitial fibrosis via TGFβ-1 pathway (56). Also, in tubular injury, Cx43 promotes ATP release and triggers macrophage pyroptosis (56, 99–101). In addition, Cx43 dysregulation in GECs disrupts NO production, promotes glycocalyx degradation, and exacerbates leakiness and oxidative stress (56, 62). Animal studies have shown that Cx43 inhibition reduces albuminuria, glomerulosclerosis, and tubular damage (56, 63, 64). Despite that, the specific mechanisms and functional consequences of Cx43 in endothelial glycolysis in DKD are still being elucidated. In this study, we examined whether barrier function is controlled via Cx43-mediated glycolytic mechanisms under diabetic conditions. Our findings showed a significant increase in Cx43 expression in diabetic mouse kidneys and confirmed robust Cx43 expression in human DKD tissues through analysis of single-nucleus RNA-seq and ATAC-seq datasets (66). In vitro, Cx43 expression was strongly induced under metabolic stress, suggesting that diabetic conditions directly regulate its expression. Mechanistically, Cx43 overexpression in endothelial cells not only enhanced glycolytic activity— evidenced by increased LDHA levels—but also disrupted VE-cadherin localization and increased endothelial permeability. Furthermore, Cx43 overexpression promoted angiogenic behavior, as seen by increased cell sprouting and cytoskeletal rearrangement. These findings establish a direct link between Cx43 and the endothelial phenotypes associated with DKD, extending the current understanding of connexins beyond intercellular communication, to include active roles in metabolic regulation and vascular integrity.

Taken together, our results suggest a novel mechanism by which diabetic conditions impair endothelial function. These insights offer new perspectives on endothelial pathophysiology in DKD and point to potential therapeutic targets. In particular, interventions aimed at suppressing pathological Cx43 activity may hold promise in preserving vascular function in diabetic patients. Finally, future studies are warranted to explore the signaling pathways upstream of Cx43 induction in diabetes, and to determine whether cell-specific Cx43 targeting (e.g., endothelial-specific knockout models) can mitigate DKD progression in vivo. Additionally, given the complex interplay between metabolism, inflammation, and vascular remodeling, it will be important to dissect how Cx43 integrates these signals to modulate endothelial fate.

## ACKNOWLEGMENTS

This work was supported partially by the National Institutes of Health (SA: R01DE031046 and R21CA294025), (ML: R01DK127830 and R01DK139676), the Toulmin Pilot Award, and by Georgetown Startup Funds to Alimperti (Assignee: 91252). This research was supported by the Microscopy and Imaging Shared Resource (MISR) and the Lombardi Comprehensive Cancer Center grant (P30-CA051008). Instrumentation used in this research included Upgrade existing multiwell Fluorescence Imaging (S10RR025661).

## AUTHOR CONTRIBUTIONS

P.L and M.L., S.A. conceptualized the idea for the study. P.L., Y.K., K.M., X.W., M.L., and S.A. designed the methodology. S.A. and M.L. acquired resources. P.L. and S.A. made the preparation for writing the original draft. P.L., Y.K., K.M., X.W., M.L., and S.A. wrote, reviewed, and edited the manuscript. All authors have read and agreed to the published version of the manuscript.

